# Comparative transcriptomics of a monocotyledonous geophyte reveals shared molecular mechanisms of underground storage organ formation

**DOI:** 10.1101/845602

**Authors:** Carrie M. Tribble, Jesús Martínez-Gómez, Fernando Alzate-Guarin, Carl J. Rothfels, Chelsea D. Specht

## Abstract

Many species from across the vascular plant tree-of-life have modified standard plant tissues into tubers, bulbs, corms, and other underground storage organs (USOs), unique innovations which allow these plants to retreat underground. Our ability to understand the developmental and evolutionary forces that shape these morphologies is limited by a lack of studies on certain USOs and plant clades. *Bomarea multiflora* (Alstroemeriaceae) is a monocot with tuberous roots, filling a key gap in our understanding of USO development. We take a comparative transcriptomics approach to characterizing the molecular mechanisms of tuberous root formation in *B. multiflora* and compare these mechanisms to those identified in other USOs across diverse plant lineages. We sequenced transcriptomes from the growing tip of four tissue types (aerial shoot, rhizome, fibrous root, and root tuber) of three individuals of *B. multiflora*. We identify differentially expressed isoforms between tuberous and non-tuberous roots and test the expression of *a priori* candidate genes implicated in underground storage in other taxa. We identify 271 genes that are differentially expressed in root tubers versus non-tuberous roots, including genes implicated in cell wall modification, defense response, and starch biosynthesis. We also identify a phosphatidylethanolamine-binding protein (PEBP), which has been implicated in tuberization signalling in other taxa and, through gene-tree analysis, place this copy in a phylogenytic context. These findings suggest that some similar molecular processes underlie the formation of underground storage structures across flowering plants despite the long evolutionary distances among taxa and non-homologous morphologies (*e*.*g*., bulbs versus tubers).

## 1 Introduction

Scientific attention in botanical fields focuses almost exclusively on aboveground organs and biomass. However, a holistic understanding of land plant evolution, morphology, and ecology requires a comprehensive understanding of belowground structures: on average 50% of an individual plant’s biomass lies beneath the ground (Niklas, 2005), and these portions of a plant are critical for resource acquisition, resource storage, and mediating the plant’s interactions with its environment. Often, belowground biomass is thought to consist solely of standard root tissue, but in some cases, plants modify “ordinary” structures for specialized underground functions. Plants called geophytes fall toward the extreme end of this belowground/aboveground allocation spectrum. In a remarkable example of convergent evolution of an innovative life history strategy, geophytes retreat underground by producing the buds of new growth on structures below the soil surface, while also storing nutrients to fuel this growth in highly modified, specialized underground storage organs (USOs) (Raunkiaer et al., 1934; Dafni et al., 1981b,a; Al-Tardeh et al., 2008; Veselý et al., 2011). Many geophytes also have the capacity to reproduce asexually through underground offshoots in addition to sexual reproduction. Geophytes are ecologically and economically important, morphologically diverse, and have evolved independently in all major groups of vascular plants except gymnosperms (Howard et al., 2019, 2020). These plants and their associated underground structures are a compelling example of evolutionary convergence; diverse taxa produce a variety of structures, often from different tissues, that serve the analogous function of underground nutrient storage. However, our understanding of the molecular processes that drive this convergence, and the extent to which these processes are themselves parallel, remains limited, due in part to the lack of molecular studies in diverse geophyte lineages. This lack of study is particularly true for monocotyledonous geophytic taxa, which comprise the majority of ecologically and economically important geophyte diversity, but have not be subject to wide scientific attention beyond a select few crops.

Some of the world’s most important crop plants have underground storage organs, including potato (stem tuber, *Solanum tuberosum*), sweet potato (tuberous root, *Ipomoea batatas*), yam (epicotyl- and hypocotyl-derived tubers, *Dioscorea* spp.), cassava (tuberous root, *Manihot esculenta*), radish (swollen hypocotyl and taproot, *Raphanus raphanistrum*), onion (bulb, *Allium cepa*), lotus (rhizome, *Nelumbo nucifera*), various *Brassica* crops including kohlrabi and turnip (Hearn et al., 2018), and more. While several of these crops are well studied and have sequenced genomes or other genetic or genomic data that may inform the molecular mechanisms underlying underground storage organ development, most detailed research has focused on a select few, which that do not represent the diversity of geophyte morphology, phylogeny, or ecology. Hearn (2006, among others) has proposed that “switches” in existing developmental programs can explain transitions between major growth forms; such a hypothesis requires broad sampling across the evolutionary breadth of taxa demonstrating the growth form. In particular, most genetic research on geophytes and their associated underground storage organs has been conducted in eudicots such as potato (Hannapel et al., 2017), sweet potato (Eserman et al., 2018; Li et al., 2019), cassava (Sojikul et al., 2010, 2015; Chaweewan and Taylor, 2015), *Brassica* (Hearn et al., 2018), and *Adenia* (Hearn, 2009). Fewer studies have focused on monocots (but see important studies in onion, such as in Lee et al., 2013), and these studies focus solely on bulbs; to date no study has characterized the molecular underpinnings of tuber formation in a monocotyledenous taxon.

This limited phylogentic breadth is particularly important in light of the findings of Hearn et al. (2018). This study provide compelling evidence that within closely related *Brassica* taxa, molecular mechanisms are shared between stem and hypocotyl/ root modifications. They also demonstrate that these mechanisms have been implicated in the development of other USOs, namely the eudicot lineages potato and sweet potato, and increased phylogenetic sampling could confirm and expand these findings or suggest that such results are clade-specific.

Underground storage organs originate from all major types of plant vegetative tissue: roots, stems, leaves, and hypocotyls. Bulbs (leaf tissue), corms (stem), rhizomes (stem), and tubers (stem or root) are some of the most common underground storage organ morphologies (Pate and Dixon, 1982), but the full breadth of morphological variation in USOs includes various root modifications (tuberous roots, taproots, etc.), swollen hypocotyls that merge with swollen root tissue (*e*.*g*., *Adenia*; Hearn, 2009), and intermediate structures such as rhizomes where the terminal end of the rhizome forms a bulb from which aerial shoots emerge (*e*.*g*., *Iris*; Wilson, 2006). Despite this morphological complexity, USOs all develop through the expansion of standard plant tissue, either derived from the root or shoot, into swollen, discrete storage organs. These storage organs also serve similar functions as belowground nutrient reserves (Veselý et al., 2011), often containing starch or other non-structural carbohydrates, storage proteins, and water. The functional and physiological similarities of underground storage organs may drive or be driven by deep molecular homology with parallel evolution in the underlying genetic architecture of storage organ development, despite differences in organismal level morphology and anatomy, as is suggested in Hearn et al. (2018).

The economic importance of some geophytes and the relevance of understanding the formation of storage organs for crop improvement have motivated studies on the genetic basis for storage organ development in select taxa. Potato has become a model system for understanding the molecular basis of USO development, and numerous studies have demonstrated the complex roles of plant hormones such as auxin, abscisic acid, cytokinin, and gibberellin on the tuber induction process (reviewed in Hannapel et al., 2017). These hormones have been additionally identified in USO formation in other tuberous root crops including sweet potato (Noh et al., 2010; Dong et al., 2019) and cassava (Melis and van Staden, 1985; Sojikul et al., 2015), in rhizome formation in *Panax japonicus* (Tang et al., 2019) and *Nelumbo nucifera* (Cheng et al., 2013b; Yang et al., 2015), and in corm formation in *Sagittaria trifolia* (Cheng et al., 2013a), suggesting that parallel processes trigger tuberization in both root- and stem-originating USOs. FT-like genes, members of the phosphatidylethanolamine-binding protein (PEBP) family, have been implicated in USO formation in potato (Navarro et al., 2011; Hannapel et al., 2017), *Dendrobium* (Wang et al., 2017), *Callerya speciosa* (Xu et al., 2016), tropical lotus (*Nelumbo nucifera*; (Yang et al., 2015), and onion (*Allium cepa*; (Lee et al., 2013), indicating either deep homology of FT involvement in USO formation across angiosperms or multiple independent involvements of FT orthologues in geophytic taxa.

The lateral expansion of roots into tuberous roots may be driven by cellular proliferation, by cellular expansion, or by a combination of these processes. Expansion in plant cells requires modification of the rigid plant cell wall to accommodate increases in cellular volume (Dolan and Davies, 2004; Humphrey et al., 2007), and genes such as expansins have been implicated in cellular expansion during tuberous root development in cassava and *Callerya speciosa* (Sojikul et al., 2015; Xu et al., 2016). Recent studies of the tuberous roots of sweet potato (*Ipomoea batatas*) and related species indicate that USO formation in these taxa involves a MADS-box gene implicated in the vascular cambium (SRD1; Noh et al., 2010) and a WUSCHEL-related homeobox gene (WOX4; Eserman et al., 2018), also involved in vascular cambium development. Additional work on cassava (*Manihot esculenta*) also suggests that tuberous root enlargement is due to secondary thickening growth originating in the vascular cambium (Chaweewan and Taylor, 2015). However, geophytes are especially common in monocotyledonous plants (Howard et al., 2019, 2020), which lack a vascular cambium entirely. No previous study has addressed the molecular mechanisms of root tuber development in this major clade, so the causes of root thickening are particularly enigmatic. Do monocots form tuberous roots through genetic machinery that shares deep homology with the eudicot vascular-cambium-related pathways, or have they evolved an entirely independent mechanism?

*Bomarea multiflora* (L. f.) Mirb. is a climbing monocotyledonous geophyte native to Venezuela, Colombia, and Ecuador, where it typically grows in moist cloud forests between 1800 *−* 3800 meters elevation (Hofreiter, 2008). *Bomarea multiflora* is an excellent model in which to study the molecular mechanisms underlying underground storage organ formation in the monocots because it has two types of underground modifications: tuberous roots and rhizomes. However, prior to this study, no genomic or transcriptomic data was available for any species of *Bomarea*. Comparative transcriptomics permits comprehensive examination of the molecular basis of development, tissue differentiation, and physiology by comparing the genes expressed in different organs, developmental stages, or ecological conditions (Ekblom and Galindo, 2011; Oppenheim et al., 2015). Because no prior genomic or transcriptomic data is needed for comparative transcriptomic studies, this method is especially appropriate for studies of non-model organisms.

In this study, we investigate the molecular mechanisms underlying the formation of tuberous roots in *Bomarea multiflora*, the first in a monocotyledonous taxon, using a comparative transcriptomics approach and quantify the extent to which these mechanisms are shared across the taxonomic and morphological breadth of geophytic taxa. Specifically, we ask by which developmental mechanisms does the plant modify fibrous roots into tuberous roots: (1) *how* expansion occurs, (2) *when* tuberization is triggered, and (3) *what* the tuberous roots store.

## 2 Materials and Methods

### 2.1 Greenhouse and Laboratory Procedures

We collected seeds from a single inflorescence of *Bomarea multiflora* in Antioquia, Colombia [vouchered as Tribble 194, deposited at UC (University Herbarium at UC Berkeley)] and germinated them in greenhouse conditions at the University of California, Berkeley designed to replicate native conditions for emphB. multiflora (70°F − 85°F and 50% humidity). Six months after germination, we harvested three sibling individuals as biological replicates. We dissected a single aerial shoot apical meristem (SAM), the rhizome apical meristem (RHI), root apical meristems (RAM) of several fibrous roots, and the growing tip of a tuberous root (TUB; Figure 1) from each of the three individuals for a total of 12 tissue samples.

**Figure 1:**
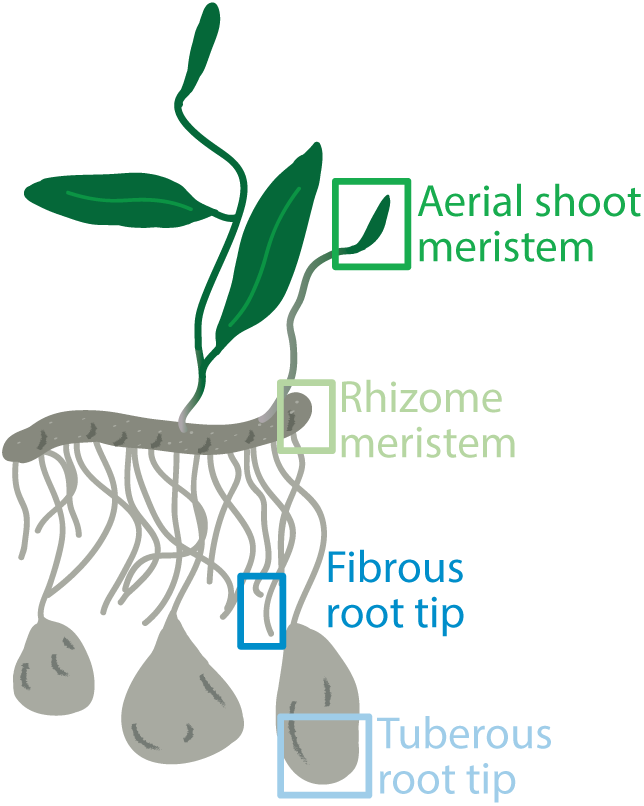
Sampling scheme of tissue types. *Bomarea multiflora* has modified underground stems (rhizomes) and modified roots (tuberous roots). We extracted RNA from the aerial shoot meristem, rhizome meristem, fibrous root tip, and tuberous root tip.

We immediately froze samples in liquid nitrogen and maintained them at *−*80^*°*^C until extraction. We extracted total RNA from all samples using the Agilent Plant RNA Isolation Mini Kit (Agilent, Santa Clara, Ca), optimized for non-standard plant tissues, especially those that may be high in starch. Quality of total RNA was measured with Qubit (ThermoFisher, Waltham MA) and Bioanalyzer 2100 (Agilent Technologies, Santa Clara, CA); if needed, we used a Sera-Mag bead clean-up to further clean extracted RNA (Yockteng et al., 2013). Two SAM samples failed to extract at sufficient concentrations, so we harvested the SAMs of two additional individuals and extracted using the Yockteng et al. (2013) protocol. Samples with an RNA integrity (RIN) score *>*7 proceeded directly to library prep. We used the KAPA Stranded mRNA-Seq Kit (Kapa Biosystems, Waltham MA) protocol for library prep, applying half reactions with an input of at least 500 ng of RNA; however, all but two samples had 1 ug of RNA. RNA fragmentation time depended on RIN score (7 *<*RIN *<*8: 4 min; 8 *<*RIN *<*9: 5 min; 9 *<*RIN: 6 min). We split samples in half after the second post-ligation clean up (Step 10 in the Kapa protocol) in order to fine-tune the enrichment step. We amplified the first half of the samples with 12 PCR cycles; this proved too low and we increased to 15 cycles for the second half of the samples. We combined samples and assessed library quality with a Bioanalyzer 2100 using the DNA 1000 kit. We performed a bead clean-up on libraries showing significant adaptor peaks (Yockteng et al., 2013). We cleaned, multiplexed, and sequenced samples on a single lane of HiSeq4000 at the California Institute for Quantitative Biosciences (QB3) Vincent J. Coates Genomics Sequencing Lab.

### 2.2 Transcriptome Assembly, Annotation, and Quantification

We cleaned, processed, and assembled the raw reads using the Trinity RNA-Seq De novo Assembly pipeline (Grabherr et al., 2011) under the default settings unless otherwise stated in associated scripts. We ran all analyses using the Savio supercomputing resource from the Berkeley Research Computing program at UC Berkeley. We cleaned reads with Trim Galore! (https://www.bioinformatics.babraham.ac.uk/projects/trim_galore/), keeping unpaired reads and using a minimum fragment length of 36 base pairs. We used all reads from all samples to generate a consensus transcriptome, assembled de novo from the concatenated data using Trinity. We aligned each sample back to the assembled consensus transcriptome using Bowtie2 (Langmead and Salzberg, 2012), and quantified isoform abundances using RSEM (RNASeq by Expectation Maximization; Li and Dewey, 2011). We assessed transcriptome assembly quality by comparing the percentages of reads that mapped back to the assembled transcriptome using Bowtie2 (Langmead and Salzberg, 2012) to align reads, among other standard metrics. We annotated the consensus transcriptome with a standard Trinotate pipeline (https://trinotate.github.io/), comparing assembled isoforms to SWISS-PROT (Boeckmann et al., 2003), RNAmmer (Lagesen et al., 2007), Pfam (Finn et al., 2014), eggNOG (Powell et al., 2014), KEGG (Tanabe and Kanehisa, 2012), and Gene Ontology (Gene Ontology Consortium, 2004) databases. We tested the concordance of biological replicates by looking for significant differences between the total number of fragments per replicate, by comparing the transcript quantities of all replicates to each other, and by checking the correlations between replicates. Isoforms with fewer than 10 total counts were discarded prior to subsequent analyses. We transformed transcript counts using the variance-stabilized transformation (VST) and compared all 12 samples using a principal components analysis.

### 2.3 Differential Expression

We identified differentially expressed isoforms (hereafter referred to as DEGs) between fibrous (FR) and tuberous (TR) roots with the R (R Core Team, 2013) DESeq2 package (Love et al., 2014). We used a p-adjusted cut-off (padj, using a Benjamini-Hochberg correction for false discovery rate) of 0.01 and a log2-fold change cut-off of 2 to determine statistically significant and sufficiently differentially expressed isoforms for downstream analyses. To test if the distribution of functional annotations for the DEGs is statistically different from the overall pool of annotated isoforms, we performed Fisher’s exact tests in R (R Core Team, 2013) to compare the distributions of number of isoforms associated with (1) biological process, (2) molecular function, and (3) cellular component GO annotations. If the overall distribution of annotations differed, we identified the specific GO terms that are enriched in the DEG dataset relative to the pool of all annotated isoforms. To identify enriched GO terms, we consider the number of isoforms associated with a particular GO category to be drawn from a binomial distribution:

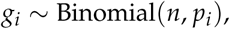

where *g*_*i*_ is the number of non-differentially expressed isoforms in GO category *i, n* is the total number of non-differentially expressed isoforms, and *p*_*i*_ is the probability that a given isoform is in GO category *i*. Thus, the maximum likelihood estimator of *p*_*i*_ is given by:

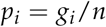

Our null hypothesis is that the probability that a given transcript is in GO category *i* is not greater in the pool of differentially expressed isoforms than the pool of non-differentially expressed isoforms. Under our null, the expected number of differentially expressed isoforms associated with GO category *i* (*k*_*i*_) is also defined by a binomial distribution:

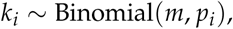

where *m* is the total number of differentially expressed isoforms and *p*_*i*_ is defined above. To test our null hypothesis, we compute the probability that the observed value is greater than *k*_*i*_, conditioning on the probability of at least one transcript count per GO category (as GO categories with zero differentially expressed transcript counts are not represented in the differentially expressed dataset) and using a Bonferroni correction (Bonferroni, 1935) for multiple comparisons.

### 2.4 Parallel Processes Across Taxa

To complement the broad survey of expression patterns, we additionally identified specific candidate genes, gene families, and molecular processes that might be involved in the development of underground storage organs via a survey of the recent literature on the molecular basis of USO formation. For each group of genes hypothesized to be involved in USO formation (either gene families or molecular/physiological processes), we first queried the annotated transcriptome for isoforms with annotations matching the associated process or family (see Supplemental Table 2 for the specific search terms used), and then tested if that group was more or less differentially expressed than expected by chance. Specifically, for each focal group of n isoforms, we randomly sampled n isoforms 10,000 times from the pool of all isoforms. For each of the 10,000 samples, we (1) compared the distribution of absolute log2-fold change values of the sampled isoforms to the distribution of log2-fold change values of the full dataset and calculated the effect size of a non-parametric Mann–Whitney–Wilcoxon test, generating an expected distribution of effect sizes for a random group of genes of size n, and (2) counted the number of significant DEGs. Under our null model, we expect the effect sizes of the focal group and the randomly sampled groups to be the same. To determine significance of the overall distributions of log2-fold change values, we compared the effect size of the focal group to the distribution of simulated effect sizes and determined if the focal group effect size fell within the 95% credible interval of the simulated distribution. Focal effect sizes that were larger than 97.5% of the simulated effect sizes indicated that the focal group is generally more differentially expressed than the null expectation; similarly, focal effect sizes smaller than 97.5% of the simulated effect sizes indicate that the focal group is less differentially expressed than the null expectation. Note that as we performed this analysis on the absolute values of log2-fold changes, less differentially expressed refers to expression levels that are more similar between the focal groups and the null than expected by chance, rather than negative log2-fold change values. Our null model also predicts that the number of significant DEGs in each focal group falls within the 95% credible set of the distributions of number of significant DEGs. We compared the number of DEGs in the focal group to distribution of number of DEGs from our 10,000 random samples. We determined significance by identifying groups that contain more or fewer DEGs than the 95% credible set of the simulated distribution.

For all targeted candidate genes, we blasted the amino acid sequence of the candidate gene to the assembled consensus transcriptome (see Supplemental Table 3 for the blasted sequence specifications) using an e-value cut-off of 0.01 to assess if the identified homologs were differentially expressed.

### 2.5 PEBP Gene Family Evolution

We reconstructed the evolutionary history of the phosphatidylethanolamine-binding protein (PEBP) gene family by combining amino acid sequences from an extensive previously published alignment (Liu et al., 2016) with the addition of sequences specifically implicated in USO formation in onion and potato or from geophytic taxa such as *Narcissus tazetta* (accession AFS50164.1), *Tulipa gesneriana* (accessions MG121853, MG121854, and MG121855), *Crocus sativa* (saffron, accession ACX53295.1), and *Lilium longiflorum* (accessions MG121858, MG121857, MG121859) (Navarro et al., 2011; Tsaftaris et al., 2012; Lee et al., 2013; Li et al., 2013; Leeggangers et al., 2017) and with copies identified in our transcriptome. For copies from *B. multiflora*, we selected the longest isoform per gene to include in the alignment. We translated coding sequences from the *Bomarea multiflora* transcriptome to amino acid sequences using TransDecoder v5.5.0 (Haas et al., 2013), removing isoforms that failed to align properly. We aligned amino acids with MAFFT as implemented in AliView v1.18 (Larsson, 2014), using trimAl v1.4.rev15 (Capella-Gutiérrez et al., 2009) with the -gappyout option. We selected the best evolutionary model with ModelTest-NG v0.1.5 (Darriba et al., 2016) (JTT+G4 amino acid substitution model (Jones et al., 1992)) and reconstructed unrooted gene trees under maximum likelihood as implemented in IQtree (Nguyen et al., 2014), run on XSEDE using the CIPRES portal (Miller et al., 2010). Using the amino acid alignment, we compared amino acid residues from *Bomarea multiflora* orthologs to those that have been functionally characterized in the *Arabidopsis* FT orthologs (reviewed in Ho and Weigel, 2014).

All scripts used in Sections 2.2, 2.3, 2.4, and 2.5 are available on GitHub at (github.com/cmt2/bomTubers)

## 3 Results

### 3.1 Transcriptome Assembly, Annotation, and Quantification

We recovered a total of 359 M paired-end 100 bp reads from the single HiSeq 4000 lane for the multiplexed 12 samples (NCBI BioProject #####). The assembled consensus transcriptome consists of 370,672 loci, corresponding to 224,661 Trinity “genes” (Dryad #####). The consensus transcriptome has a GC content of 45.14%, N50 of 1191 bp, median transcript length of 317 bp, and mean transcript length of 556.95 bp (see also Supplemental Materials Section 1.1). Of all reads, 85.54% aligned concordantly (in a way which matches Bowtie2’s (Langmead and Salzberg, 2012) expectation for paired-end reads) to the assembled transcriptome, indicating sufficient assembly quality to proceed with downstream analyses (see Supplemental Materials section 1.2). 8.15% of genes had at least 10% sequence identity with the UniProt database (See Supplemental Material section 1.3, Table 1 and Figure 1; Consortium, 2019). Of those, 53.49% blasted with at least 80% sequence identity. The Trinotate pipeline annotated 1.70% of all isoforms. All four tissue types showed concordance between the three biological replicates with generally 1:1 ratios of transcript quantities to each other (see Supplemental Materials section 2: Figures 2 - 5) so we proceeded with analyses using data from all three biological replicates.

**Table 1:** Top ten most differentially expressed isoforms (with *padj <* 0.01) and their corresponding annotations.

| Transcript ID: Annotated Name (SPROT) | Log2 Fold Change | Gene Ontology |  |  |
| --- | --- | --- | --- | --- |
|  |  | Cellular Components | Molecular Functions | Biological Processes |
| TRINITY_DN116220.c0.g1.i: Zinc finger CCCH domain-containing protein 55 | 40.25 |  | DNA binding, metal ion binding |  |
| TRINITY_DN128685.c1.g3.i4: Callose synthase 3 | 33.85 | 1,3-beta-D-glucan synthase complex, integral component of membrane, plasma membrane | 1,3-beta-D-glucan synthase activity | (1-3)-beta-D-glucan biosynthetic process, cell wall organization, regulation of cell shape |
| TRINITY_DN121298.c2.g2.i: Heat shock 70 kDa protein 15 | 33.52 | cell wall, cytosol, nucleus, plasma membrane, plasmodesma | ATP binding |  |
| TRINITY_DN115892.c0.g1.i3: Pre-mRNA-splicing factor ATP-dependent RNA helicase DEAH1 | 33.52 | membrane, spliceosomal complex | ATP binding, ATP-dependent 3'-5' RNA helicase activity, RNA binding | mRNA processing, posttranscriptional gene silencing by RNA, RNA splicing |
| TRINITY_DN127064.c0.g3.i: LRR receptor-like serine/threonine-protein kinase HSL2 | 33.32 | integral component of membrane, plasma membrane | ATP binding, protein serine/threonine kinase activity | defense response to Gram-negative bacterium, lateral root morphogenesis, leaf abscission, regulation of gene expression |
| TRINITY_DN128839.c3.g1.i6: Palmitoyl-acyl carrier protein thioesterase, chloroplastic | 33.01 | chloroplast | thiolester hydrolase activity | fatty acid biosynthetic process |
| TRINITY_DN121430.c10.g2: Sucrose nonfermenting 4-like protein | 32.64 | chloroplast, chloroplast starch grain, cytoplasm, nucleus | kinase binding, maltose binding, protein kinase activator activity, protein kinase regulator activity, protein serine/threonine kinase activity | carbohydrate metabolic process, cellular response to glucose starvation, mitochondrial fission, peroxisome fission, pollen hydration, protein autophosphorylation, regulation of protein kinase activity, regulation of reactive oxygen species metabolic process |
| TRINITY_DN124527.c1.g1.i5: Bifunctional TH2 protein, mitochondrial | 32.64 | cytosol, mitochondrion | thiaminase activity, thiamine phosphate phosphatase activity | thiamine biosynthetic process |
| TRINITY_DN120224.c4.g1.i: Enhancer of polycomb homolog 2 | 32.25 | Piccolo NuA4 histone acetyltransferase complex |  | DNA repair, histone acetylation, regulation of transcription by RNA polymerase II |
| TRINITY_DN122787.c0.g1.i1: Protein IQ-DOMAIN 32 | -32.16 | chloroplast envelope, cytosol, microtubule associated complex, nucleus, plasma membrane |  | response to abscisic acid |

**Figure 2:**
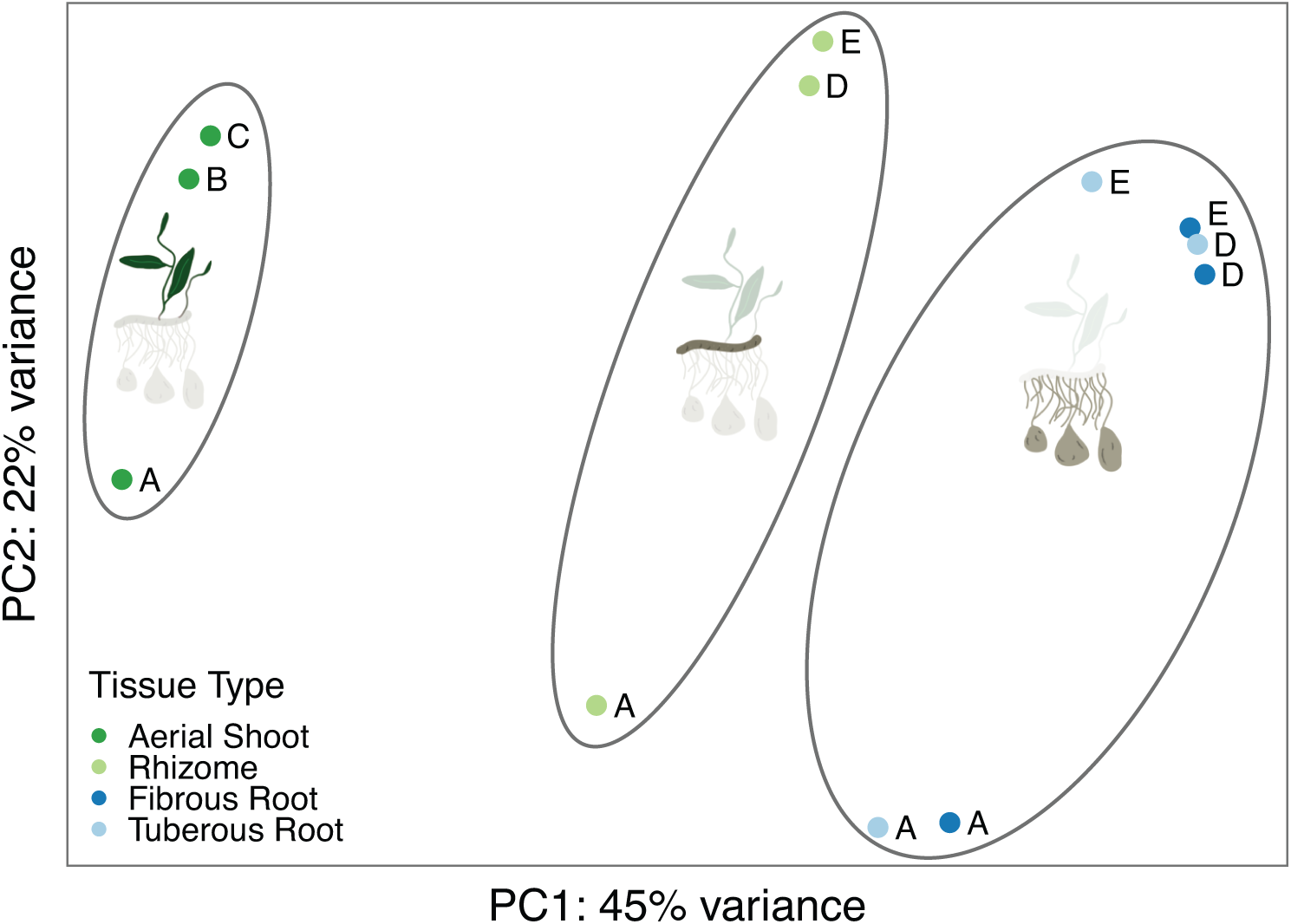
Principal component analysis of VST-transformed transcript counts from all samples. We performed a principal component analysis of variance-stabilizing-transformed (VST) transcript counts from all biological replicates of all tissue types. Points are colored by tissue type, and letters correspond to the individual plants sampled.

A principal component analysis (PCA) of the VST transcript counts (Figure 2) shows that the first PC axis (45% of the variance in samples) generally explains the variation between tissue types. The shoot tissues (SAM and RHI) cluster separately, while the root tissues (ROO and TUB) cluster together. The underground rhizome samples (RHI) fall out intermediate between the aerial shoot samples (SAM) and the underground root and tuber samples (ROO and TUB) along this axis. The co-clustering of fibrous and tuberous root samples in the PCA indicates that the overwhelming, general pattern of expression in all root samples is similar, especially in contrast to the very distinct expression profiles of the shoot samples. The second PC axis (22% of variance) generally explains variance among biological replicates, with Individual A particularly distinct from other individuals, perhaps due to microhabitat variation in the greenhouse, genotype differences, increased or decreased herbivory compared to other individuals.

### 3.2 Differential Expression

We recovered a total of 271 differentially expressed isoforms (DEGs) between fibrous and tuberous roots (FR vs. TR). Of these, 226 correspond to regions of the assembled consensus transcriptome with functional annotations.

Of the three types of Gene Ontology (GO) annotations, we recovered significant differences in the distributions of the number of isoforms associated with GO categories between the differentially expressed dataset and the pool of all isoforms for molecular functions (*p* = 1*x*10^*−*4^) and cellular components (*p* = 0.031) but not for biological processes (*p* = 0.921). We recovered no significantly enriched individual GO annotations for biological processes. For cellular components, we found that *cytoplasm, integral component of membrane, nucleus*, and *plasma membrane* were enriched in the differentially expressed dataset relative to non-differentially expressed isoforms. For molecular functions, we found that *ATP binding* and *metal ion binding* were enriched.

Of the 271 DEGs, 126 (46.5%) were over-expressed in tuberous roots while the remaining 145 (53.5%) were under-expressed. All top ten most differentially expressed isoforms (the ten DEGs with the highest absolute value log2-fold change values between fibrous and tuberous roots) are implicated in cellular and biological processes (Table 1). All but one of these top ten DEGs are overexpressed in tuberous roots and are generally implicated in nucleotide and ATP binding, cell wall modification, root morphogenesis, and carbohydrate and fatty acid biosynthesis. The most differentially expressed isoform (with a 40.25 log2-fold change), TRINITY DN116220 c0 g1 i4, is a Zinc finger CCCH domain-containing protein 55, a possible transcription factor of unknown function. Other notable top DEGs include TRINITY DN128685 c1 g3 i4, callose synthase 3, which regulates cell shape, TRINITY DN121298 c2 g2 i5, a heat shock protein, TRINITY DN127064 c0 g3 i1, an LRR receptor-like serine implicated in lateral root morphogenesis, and TRINITY DN121430 c10 g2 i, a carbohydrate metabolism protein. The tenth most differentially expressed DEG, under-expressed in tuberous roots, is implicated in abscisic acid signaling.

### 3.3 Parallel Processes Across Taxa

Based on our literature survey, we identified 12 groups of genes that have been implicated in USO formation in other geophytes: abscisic acid response genes, calcium-dependent protein kinases (CDPK), expansins, lignin biosynthesis, MADS-Box genes, starch biosynthesis, auxin response genes, cytokinin response genes, 14-3-3 genes, gibberellin response genes, KNOX genes, and lipoxygenases (See Figure 3 and Supplemental Table 4). Of these 12 gene groups, six have significantly different expression values than the overall distribution of expression for all isoforms. In four cases (expansins, lignin, MADS-Box, and starch biosynthesis) the overall expression levels are significantly more differentially expressed than expected by chance. Expansins and lignin are generally under-expressed in TR compared to FR while MADS-Box and starch are generally over-expressed in TR compared to FR. In two cases (14-3-3 and lipoxygenases), the overall expression levels are less differentially expressed than expected by chance; their expression levels are generally conserved between TR and FR. In the remaining six cases (abscisic acid, auxin, CDPK, cytokinin, gibberellin, and knox) we recover no significant pattern in overall expression levels.

**Figure 3:**
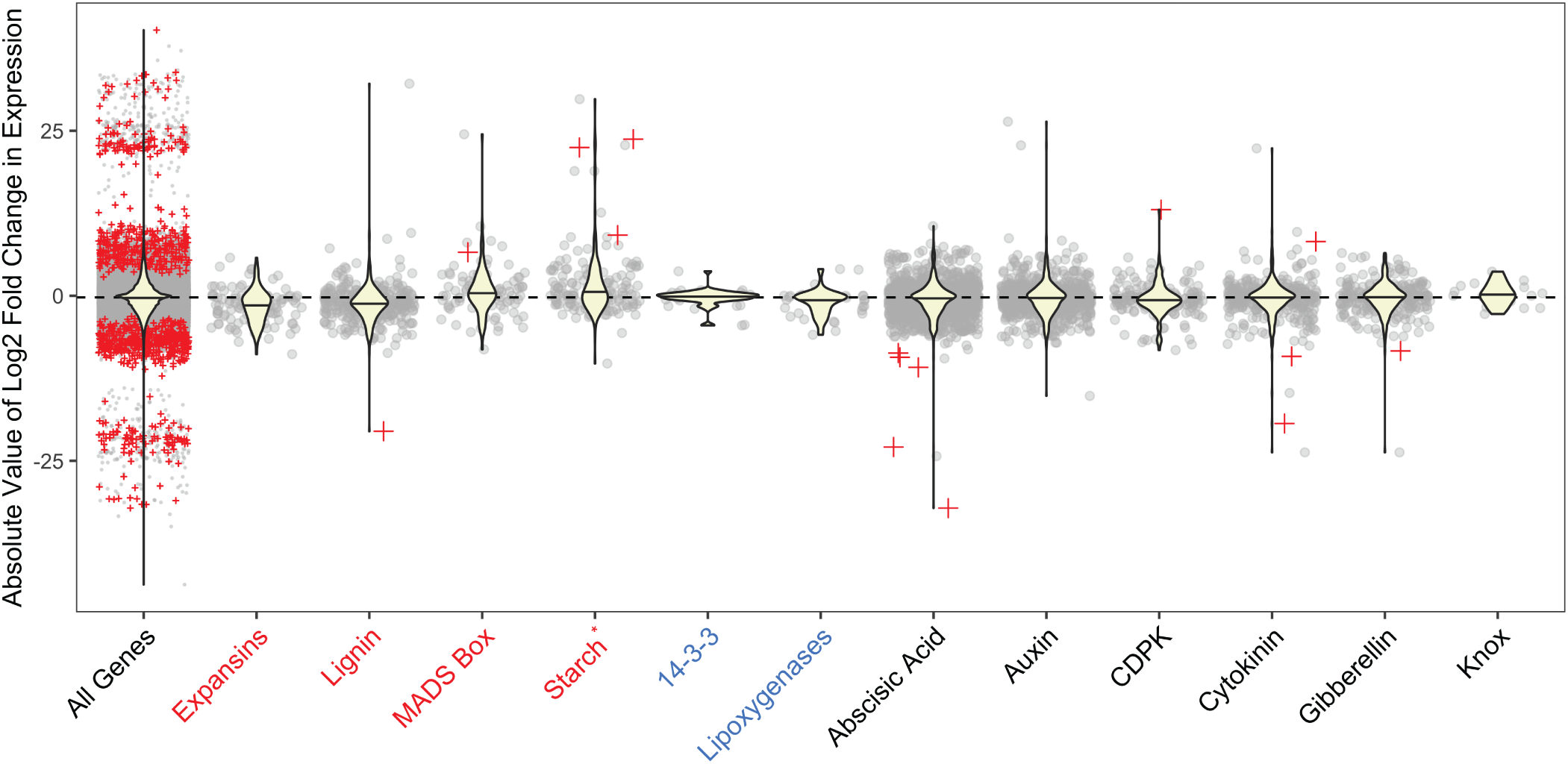
Differential expression of candidate gene groups. We identified isoforms corresponding to specific pathways and gene families (groups of genes) and categorize the log2-fold change of expression between tuberous and fibrous roots of those groups. Positive values correspond to overexpression in tuberous vs. fibrous roots. Box plots correspond to the log2-fold change value for the gene groups and grey points correspond to individual isoforms. Isoforms that are significantly differentially expressed (padj *<*0.01) are labeled as red asterisks within the scatter plots. Groups with absolute value log2-fold change distributions that are significantly larger from the entire dataset (shown in All Genes) are labeled in red on the X-axis, indicating that these groups are more differentially expressed (either generally up or down in tuberous roots) than expected:expansins, lignin, MADS box, and starch. Groups with less differential expression than expected by chance are labeled in blue: 14-3-3 and lipoxygenases. Groups with significantly more differentially expressed isoforms that expected by chance are labeled with an asterisk on the X-axis: starch.

We identified fifteen individual DEGs in these gene groups (Table 2); interestingly, there seems to be no generalizable relationships between the significance and directionality of a particular group’s distribution with the presence and directionality of expression for individual DEGs. For example, the expression distribution of gibberellin genes does not deviate significantly from the global pool of isoforms, but there is one significantly under-expressed DEG in the gibberellin group; similarly, the CDPK genes are under-expressed as a group, but the only CDPK-related significant DEG is over-expressed (Table 2).

**Table 2:**
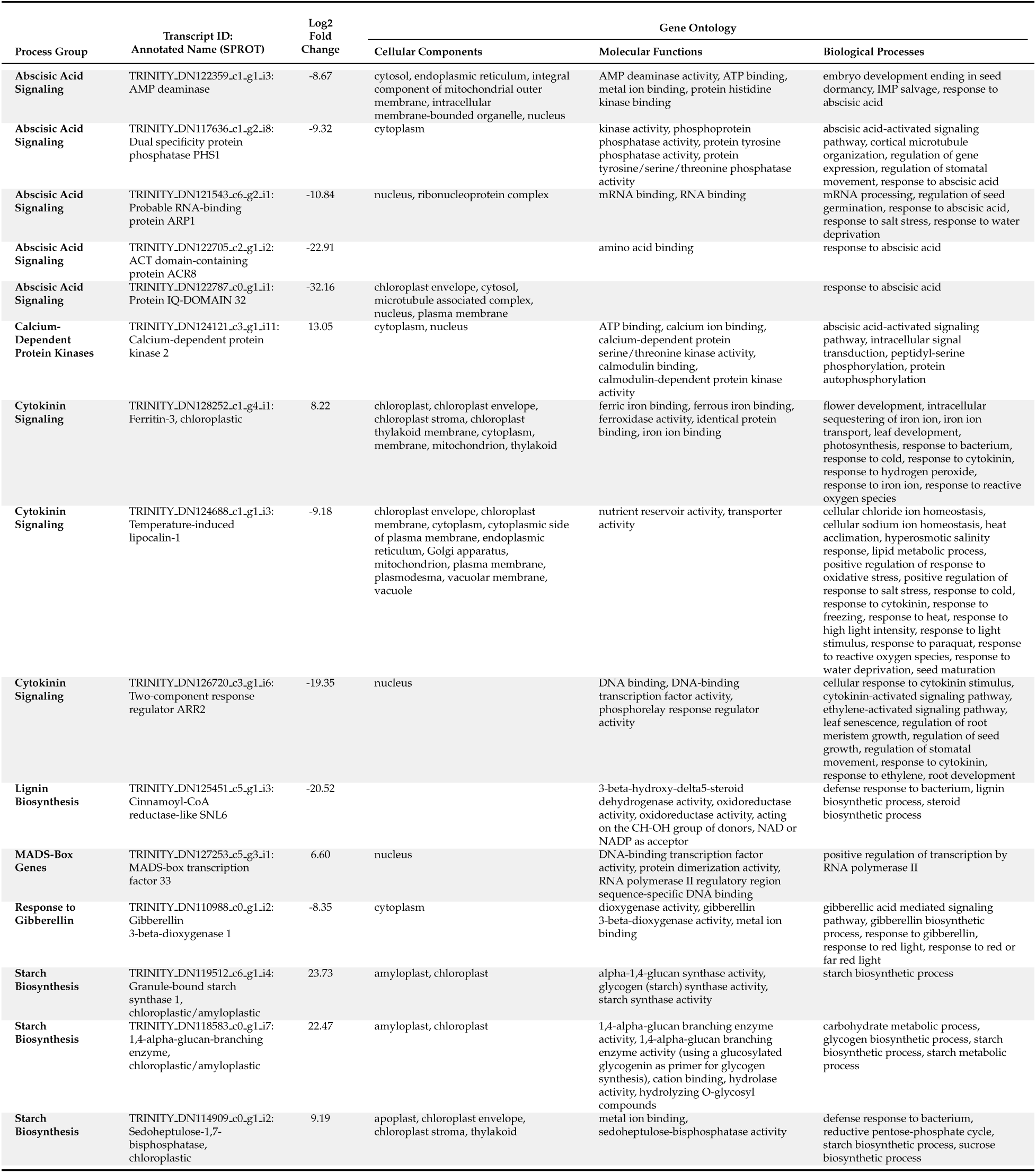
Differentially expressed isoforms (with *padj <* 0.01) in specific gene groups and their corresponding annotations.

In addition to the gene groups, we identified five specific candidate genes from the literature: OsbHLH120 (qRT9) has been implicated in root thickening in rice (Li et al., 2015); IDD5 and WOX4 are implicated in starch biosynthesis and TR formation, respectively, in Convolvulaceae (Eserman et al., 2018); sulfite reductase is associated with TR formation in *Manihot esculenta* (Sojikul et al., 2010); and FLOWERING LOCUS T (FT) has been implicated in signaling the timing of USO formation in a variety of taxa, notably *Allium cepa* and *Solanum tuberosum* (Navarro et al., 2011; Hannapel et al., 2017). We recover between nine and 63 putative homologs of these candidates (Table 3), but only one is significantly differentially expressed (*padj <* 0.01): a putative FT homolog (TRINITY DN129076 c1 g1 i1), further investigated in the PEBP Gene Family Evolution section (below). One putative qRT9 homolog is marginally significant (*padj* = 0.050), and the E-value from the BLAST result to this isoform was 0.09. Given these marginal significance values, it is likely that the result is spurious and we do not follow up with further analysis.

**Table 3:** Specific candidate genes and results from blasting to assembled transcriptome. Asterisk indicates marginal significance. Isoforms found corresponds to the number of copies identified in the assembled *Bomarea multiflora* transcriptome, and Number DE corresponds to the number of those copies which are significantly differentially expressed.

| Gene Names | Original Taxon | Isoforms Found | Number DE |
| --- | --- | --- | --- |
| OsbHLH120 | Oryza sativa | 22 | 1* |
| IDD5 | Ipomea batatas | 63 | 0 |
| WOX4 | Ipomea batatas | 31 | 0 |
| Sulfite reductase | Manihot esculenta | 9 | 0 |
| FT-like | Solanum tuberosum, Allium cepa | 37 | 1 |

### 3.4 PEBP Gene Family Evolution

We identified thirty-seven *Bomarea* isoforms as putative FLOWERING LOCUS T (FT) homologs. After filtering for the longest isoform per gene and filtering out sequences which failed to align properly, ultimately, we include five sequences in addition to the significantly differentially expressed copy. We recover three major clusters in our unrooted gene tree, all with strong bootstrap support (Figure 4a); these correspond to the FT cluster, TERMINAL FLOWER 1 (TFL1) cluster, and MOTHER OF FT AND TFL1 (MFT) cluster recovered in previous analyses (Liu et al., 2016). Three of the six *Bomarea multiflora* isoforms fall out in the FT cluster and three in the TFL1 cluster. The *Bomarea* DEG homolog is highly supported in the TFL1 cluster with sequences from other monocot taxa (Figure 4b). Members of this TFL1 clade have been functionally characterized in *Oryza sativa*, where four orthologs (OsRNC1, OsRNC2, OsRNC3 and OsRNC14) antagonize the rice ortholog of FT to regulate inflorescence development (Kaneko-Suzuki et al., 2018). mRNA isoforms of all OsRNC are expressed in the root and transported to the SAM. Interestingly, the Crocus sativus ortholog (CsatCEN/TFL1) belongs to the same clade and is also expressed underground (in corms; Tsaftaris et al., 2012). Amino acid analysis of TRINITY DN129076 c1 g1 i1.p1 shows that it convergently shares a glycine residue at AthFT position G137, typically characteristic of FT rather than TFL genes (Supplemental Figure 8 Pin et al., 2010). FT homologs from *Allium cepa* and *Solanum tuberosum* that have been functionally implicated in stem tuber and bulb formation, respectively, are in the FT cluster but do not cluster together; rather all USO-implicated PEBP genes are more closely related to non-USO copies than to each other.

**Figure 4:**
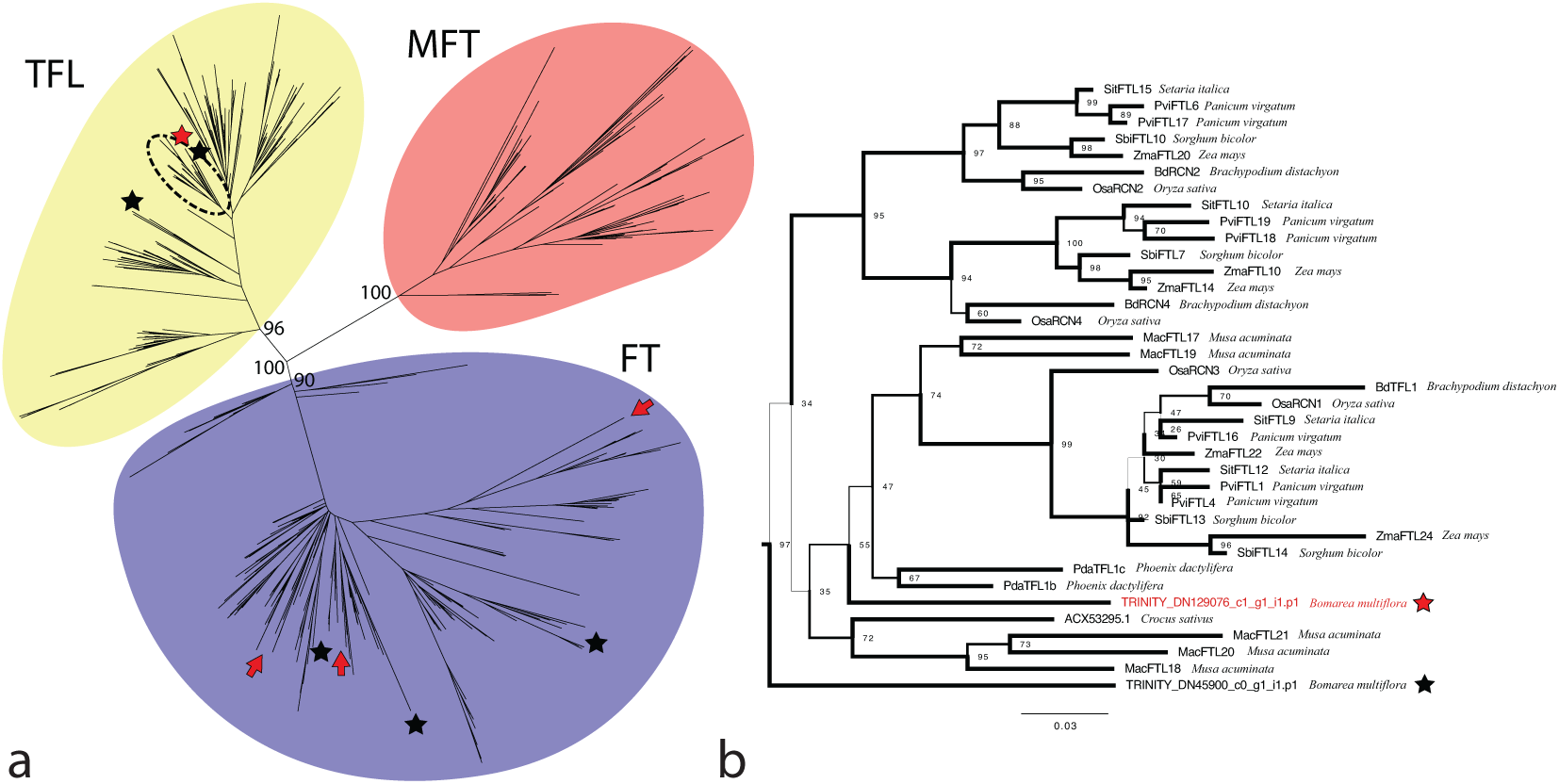
Evolution of PEBP genes: (a) Unrooted gene tree of 540 PEBP gene copies from across land plants. Stars indicate all included copies from *Bomarea multiflora*; the red star corresponds to the significantly differentially expressed isoform TRINITY DN129076 c1 g1 i1). Red and arrows indicate PEBP copies that have been implicated in USO formation in other taxa (*Allium cepa* and *Solanum tuberosum*). The major clusters correspond to the FLOWERING LOCUS T (FT, in purple), TERMINAL FLOWER 1 (TFL1, in yellow), and the MOTHER OF FLOWERING LOCUS T AND TERMINAL FLOWER 1 (MFT, in red) gene groups and are labeled with high bootstrap support. (b) Detailed view of the cluster indicated by a dashed circle in (a), including monocot-specific copies of TERMINAL FLOWER 1 genes. Line thickness corresponds to bootstrap support.)

## 4 Discussion

### 4.1 How to Make a Tuberous Root

Our results suggest potential developmental mechanisms by which the plant modifies fibrous roots into tuberous roots: 1) *how* expansion occurs, 2) *when* tuberization is triggered, and 3) *what* the tuberous roots store.

Root expansion likely occurs due to primary thickening growth via cellular expansion *B. multiflora*. Due to the absence of a vascular cambium, secondary growth is not likely to be involved, despite the prevalence of this mechanism in other taxa such as sweet potato (Noh et al., 2010; Eserman et al., 2018) and cassava (Melis and van Staden, 1985).

Cell wall-related genes include pectinesterase TRINITY DN122210 c6 g1 i1 (log2-fold change = 21.91; padj = 1.22E-8), which modifies pectin in cell walls leading to cell wall softening, as demonstrated for example in *Arabidopsis* (Braybrook and Peaucelle, 2013). Interestingly, expansins as a group were under-rather than over over-expressed in *B. multiflora*, though this is the mechanism by which cell expansion occurs in other taxa (see Expansins discussion below). Two of the enriched cellular component GO categories involve modifications to the cell membrane (*integral component of membrane* and *plasma membrane*), which suggests that modifications to the membrane may also be necessary in cellular expansion.

Flowering development genes such as TFL genes may be involved in mediating environmental signals and inducing tuber formation. Tuberization signaling may also be mediated by callose production, influencing symplastic signaling pathways through plasmodesmata modification. Callose synthase 3 is one of the most highly differentially expressed DEGs (TRINITY DN128685 c1 g3 i4, Table 1). Callose is a much less common component of cell walls than cellulose (Schneider et al., 2016), but it is often implicated in specialized cell walls and in root-specific expression (Vatén et al., 2011; Benitez-Alfonso et al., 2013). Callose synthase has been implicated in the development of other unique root-based structures such as root nodules (Gaudioso-Pedraza et al., 2018) and mutations in callose synthase 3 affect root morphology (Vatén et al., 2011), suggesting that callose synthase 3 may play an integral role in triggering tuberous roots development in *B. multiflora* through symplastic signaling pathways and/or in modifying cell walls to accomodate expansion. Callose involvment in USO formation has not previously been reported and may be unique to *B. multiflora* or to monocots.

Finally, starch is thought to be the primary nutrient reserve in *Bomarea* tubers (Kubitzki and Huber, 1998). Many previous studies have found evidence of overexpression of carbohydrate and starch synthesis molecules in USOs (for example in sweet potato; Eserman et al., 2018). Differentially expressed isoforms implicated in the carbohydrate metabolic process support the presence of active starch synthesis in our data. One of the most differentially expressed isoforms is a homolog of sucrose non-fermenting 4-like protein (Table 1, TRINITY DN121430 c10 g2 i1) and participates in carbohydrate biosynthesis, demonstrating that *B. multiflora* tubers were actively synthesizing starch when harvested. Additionally, genes implicated in defense response, such as TRINITY DN127064 c0 g3 i1 (LRR receptor-like serine/ threonine-protein kinase HSL2, Table 1) may be differentially expressed in tuberous roots to protect starch reserves against potential predation by belowground herbivores. LRR receptors have been implicated in triggering various downstream plant immune responses (Liang and Zhou, 2018).

We emphasize that more detailed work to follow up on these aspects of root tuber development should include morphological, anatomical, and developmental characterization of the tuberization process. The integration of these methods with genetic and molecular characterization will likely provide important functional links between the observed expression patterns we report here and the specific effects on plant form and function.

### 4.2 Similarities in Molecular Mechanisms of USO Formation

We identify four molecular processes, previously implicated in USO formation in other taxa, which are either over- or under-expressed in the tuberous roots of *Bomarea multiflora* (Figure 3). These processes suggest potential parallel function across deeply divergent evolutionary distances and in distinct plant structures.

***Expansins*** are cell wall modifying genes known to loosen cell walls in organ formation (Dolan and Davies, 2004; Humphrey et al., 2007; Braybrook and Peaucelle, 2013). Their involvement in USO formation has been documented in the tuberous roots of cassava (Sojikul et al., 2015) and *Callerya speciosa* (Xu et al., 2016), the rhizomes of *Nelumbo nucifera* (Cheng et al., 2013b), the tuberous roots of various Convolvulaceae (Eserman et al., 2018), and the stem tubers of potato (Jung et al., 2010). As a group, in our data expansins are under-expressed in tuberous compared to fibrous roots, but no individual DEGs are statistically significant. These results suggest, surprisingly, that expansins likely do not play an important role in tuberous root formation in *Bomarea multiflora*. We do identify callose synthase 3 as one of the most differentially expressed genes (Table 1, so it is possible that cell wall modification during tuber development is primarily driven by callose rather expansin action in *B. multiflora*.

***Lignin biosynthesis genes*** are under-expressed in several geophytic taxa with tuberous roots, including cassava (Sojikul et al., 2015), wild sweet potato (*Ipomoea trifida*; (Li et al., 2019), and *Callerya speciosa* (Xu et al., 2016). Similarly, we find that lignin biosynthesis overall is under-expressed in tuberous compared to fibrous roots, and one isoform in particular is significantly underexpressed: TRINITY DN125451 c5 g1 i3: Cinnamoyl-CoA reductase-like SNL6 (Table 2). This gene has been found to significantly decrease lignin content without otherwise affecting development in tobacco (Chabannes et al., 2001). Decreased lignin in tuberous roots may further allow for cell expansion and permit lateral swelling of tuberous roots during development.

***MADS-Box genes*** are implicated in USO formation in the tuberous roots of wild sweet potato (*Ipomoea trifida* (Li et al., 2019) and sweet potato (*Ipomoea batatas* (Noh et al., 2010; Dong et al., 2019), the rhizomes of *Nelumbo nucifera* (Cheng et al., 2013b), and the corms of *Sagittaria trifolia* (Cheng et al., 2013a), indicating widespread parallel use of MADS-Box genes in the formation of USOs. Similarly, we find that MADS-Box genes overall, and one DEG in particular, are over-expressed in *Bomarea* tuberous roots. MADS-Box genes are implicated widely as important transcription factors regulating plant development (Buylla et al., 2000). It is thus unsurprising that MADS-Box genes are regularly implicated in USO formation. It remains unclear if the MADS-Box genes identified in the aforementioned studies represent independent involvement of MADS-Box genes in USOs from other aspects of plant development, or if they form a clade of USO-specific copies. Follow-up phylogenetic analyses of these genes could prove informative in determining if MADS-Box genes involved in USO development form a clade (perhaps indicating a common origin for MADS-Box involvement in USOs or molecular convergence) or if they fall out independently (perhaps indicating multiple events of MADS-box involvement through distinct molecular changes).

***Starch*** biosynthesis genes are very commonly identified in the formation of USOs, including in cassava (Sojikul et al., 2010, 2015), *Nelumbo nucifera* (Cheng et al., 2013b; Yang et al., 2015), wild and domesticated sweet potatoes (Eserman et al., 2018; Li et al., 2019; Dong et al., 2019), and potato (Xu et al., 1998). Since starch is so ubiquitous in USOs, this is unsurprising. We also find starch isoforms overall to be over-expressed in *Bomarea* tuberous roots, and three genes in particular are significantly over-expressed (see Table 2).

Unexpectedly, two of our candidate gene groups, 14-3-3 genes and lipoxygenases, show significantly less differential expression that expected by chance. This result suggests that some groups of genes previously identified as involved in USO formation may be conserved in their expression patterns between tuberous and fibrous roots in *Bomarea multiflora* unlike their differential expression patterns in other taxa. The evolutionary origin of 14-3-3 and lipoxygenase gene families likely pre-dates the divergence of plant groups and the evolution of any kind of USO (both gene families are found across eukaryotes), implying that their involvement in USO development is perhaps due to neofunctionalization in specific plant clade(s). To our knowledge, the gene families’ involvement in USO development has so far been only documented in eudicots, suggesting that this neofunctionalization may have occurred in the eudicot clade and would thus not be documented in monocots, a counter-example to parallelism in USO development across land plants. Gene tree analysis of these families and the homologs implicated in USO development would yield further insight into these potential patterns.

The other molecular processes we tested failed to show group-level differences from the global distribution of expression levels. However, the presence of DEGs in some of these groups indicates that the phytohormones in particular may play a role in tuberous root formation. One gibberellin response isoform is significantly under-expressed in tuberous roots, which aligns with previous research suggesting that decreased gibberellin concentrations in roots can lead to root enlargement (Tanimoto, 2012) and tuber formation (Xu et al., 2016; Li et al., 2019; Dong et al., 2019). The lack of significant auxin-related isoforms as differentially expressed is surprising, as auxin has been implicated in USO formation in several previous studies (Noh et al., 2010; Cheng et al., 2013b; Sojikul et al., 2015; Yang et al., 2015; Xu et al., 2016; Hannapel et al., 2017; Li et al., 2019; Dong et al., 2019; Kolachevskaya et al., 2019), but it is possible this paritcular role of auxin is not part of tuberous root development in monocot taxa, or that it is simply not identified in this study.

### 4.3 PEBP and FT-Like Gene Evolution in Geophytic Taxa

Gene tree analysis of PEBPs indicates that copies of these genes have independently evolved several times to signal USO formation in diverse angiosperms (including monocots and eudicots) and in diverse USO morphologies (including tuberous roots, bulbs, and stem tubers). Furthermore, the presence of TFL1 and FT homologs in gymnosperms and other non-flowering plants (Liu et al., 2016) suggests that the origin of these genes predates the evolution of the flower, their name notwithstanding. Instead, it seems likely that these genes originally evolved as environmental signaling genes with wider involvement in triggering the seasonality of various aspects of plant development. Subsequently, it is possible that gene duplication followed by neofunctionalization caused many copies in flowering plants to signal flower develepment and other copies to signal USO development. Given our results, it is possible that USO-specialized FT and TFL1 genes arose at least four times independently, suggesting that broadly parallel molecular evolution may underlie the convergent morphological evolution of USOs. However, without further work to quantify and characterize the differentially expressed PEBP gene in *Bomarea multiflora*, it is unclear if this gene is involved in tuberization signalling. Previously verified USO-specialized PEBP genes are clearly FT genes, but the *Bomarea multiflora* copy is a TFL1 ortholog. In the *Beta vulgaris* FT ortholog BvFT1, a mutation from glycine to glutamine along with two other mutations are sufficient to turn FT into an antagonist of the functional FT orthology BvFT2 (Pin et al., 2010). None of the other TFL1 orthologs in the monocot clade share this mutation. While the *B. multiflora* ortholog does show signs of molecular convergence with FT genes at one residue known to induce TFL-like function in FT genes 8, it is unknown if function recovery can occur from TFL to FT and if only one mutation out of three is sufficient to recover function. Follow up studies to characterize the functions of various TFL1 and FT orthologs in *Bomarea multiflora* are necessary understand the role this DEG plays in tuber development.

Furthermore, the identification of additional USO-specific PEBP genes would shed more light on patterns of PEBP family involvement in USOs, but the dearth of studies on the molecular basis of USO development impedes such analyes. With increased sampling, followup studies could identify unique patterns of convergent molecular evolution on the USO-specific FT genes, such as tests for selection or further characterization of independently derived subsequences or motifs that could reflect or cause shared function.

### 4.4 Conclusions

We provide the first evidence of the molecular mechanisms of tuberous root formation in a monocotyledonous taxon, filling a key gap in understanding the commonalities of storage organ formation across taxa. We demonstrate that several groups of genes shared across geophytic taxa are implicated in tuberous root development in *Bomarea multiflora*, patterns which suggest that deep parallel evolution at the molecular level may underlie the convergent evolution of an adaptive trait. In particular, we demonstrate that PEBP genes previously implicated in underground storage organ formation appear multiple times across the gene tree, and we suggest that a TFL1 gene in *Bomarea multiflora* may also be involved in USO development. These patterns imply that repeated morphological convergence may be matched by independent evolutions of similar molecular mechanisms. However, we also demonstrate two counter examples to this pattern: groups of genes previously implicated in USO development whose patterns of expression are more similar in tuberous and fibrous roots than expected by chance (14-3-3 genes and lipoxygenases). These findings suggest further avenues for research on the molecular mechanisms of how plants retreat underground and evolve strategies enabling adaptation to environmental stresses. More molecular studies on diverse, non-model taxa and more thorough sampling of underground morphological diversity will enhance our understanding of the full extent of these convergences and add to our general understanding of the molecular basis for adaptive, convergent traits.

## 4.5 Acknowledgements

This research used the Savio computational cluster resource provided by the Berkeley Research Computing program at the University of California, Berkeley (supported by the UC Berkeley Chancellor, Vice Chancellor for Research, and Chief Information Officer). We additionally thank Lydia Smith (Evolutionary Genetics Lab, UC Berkeley) for training and sharing her expertise on RNA-Seq, NSF GRFP, SSB, ASPT, Pacific Bulb Society, UC Berkeley’s Integrative Biology Department, and the Tinker Foundation for support to CMT, and UC Berkeley CNR and the University and Jepson Herbaria for supporting sequencing costs. Michael R. May, David D. Ackerly, and Benjamin K. Blackman, and two anonymous reviewers provided feedback on the manuscript.

