## Supplemental Materials for "Comparative transcriptomics of a monocotyledonous geophyte reveals shared molecular mechanisms of underground storage organ formation"

### Supplemental Material

#### 1 Quality checks of transcriptome assembly

##### 1.1 Basic statistics using Trinity's TrinityStats.pl

- Counts of transcripts, etc.
  - Total trinity 'genes': 224661
  - Total trinity transcripts: 370672
  - Percent GC: 45.14
- Stats based on ALL transcript contigs
  - Contig N10: 3633
  - Contig N20: 2661
  - Contig N30: 2059
  - Contig N40: 1595
  - Contig N50: 1191
  - Median contig length: 361
  - Average contig: 691.60
  - Total assembled bases: 256357337
- Stats based on ONLY LONGEST ISOFORM per 'GENE'
  - Contig N10: 3246
  - Contig N20: 2280
  - Contig N30: 1656
  - Contig N40: 1149
  - Contig N50: 763
  - Median contig length: 317
  - Average contig: 556.95
  - Total assembled bases: 125124081

These statistics are in line with the expectations for de-novo transcriptome assembly using the Trinity Pipeline (see, for example: `https://github.com/trinityrnaseq/trinityrnaseq/wiki/There-are-too-many-transcripts!-What-do-I-do%3F`. It is unlikely that the high number of Trinity 'genes' is due to allelic variation, as Trinity accounts for potential allelic variants in the assembly process (Grabherr et al., 2011). In addition, *Bomarea* and monocots in general tend to have large genomes, which may also explain the high number of trinity 'genes'.

#### 1.2 Assessing the Read Content of the Transcriptome Assembly

From tutorial: `https://github.com/trinityrnaseq/trinityrnaseq/wiki/RNA-Seq-Read-Representation-by-Trinity-Assembly`

- 324348568 reads; of these:
  - 324348568 (100.00%) were paired; of these:
    - 46883155 (14.45%) aligned concordantly 0 times
    - 73867485 (22.77%) aligned concordantly exactly 1 time
    - 203597928 (62.77%) aligned concordantly >1 times

---

  - 46883155 pairs aligned concordantly 0 times; of these:
    - 1219749 (2.60%) aligned discordantly 1 time

---

  - 45663406 pairs aligned 0 times concordantly or discordantly; of these:
    - 91326812 mates make up the pairs; of these:
      - 45313307 (49.62%) aligned 0 times
      - 10083388 (11.04%) aligned exactly 1 time
      - 35930117 (39.34%) aligned >1 times

##### 1.3 Full-length transcript analysis for model and non-model organisms using BLAST+

From tutorial: `https://github.com/trinityrnaseq/trinityrnaseq/wiki/Counting-Full-Length-Trinity-Transcripts`

| **hit_pct_cov_bin** | **count_in_bin** | **> bin_below** |
|---|---|---|
| 100 | 6217 | 6217 |
| 90 | 2210 | 8427 |
| 80 | 1368 | 9795 |
| 70 | 1102 | 10897 |
| 60 | 1142 | 12039 |
| 50 | 1209 | 13248 |
| 40 | 1344 | 14592 |
| 30 | 1623 | 16215 |
| 20 | 1592 | 17807 |
| 10 | 504 | 18311 |

**Table 1:** Results of blasting transcriptome 'gene's to the UniProt database (Consortium, 2019). count.in.bin shows the number of genes that have a certain interval (e.g. 70-80%) sequence identity with the blast hit, >bin.below shows the number of genes with at least than interval (e.g. at least 70%)

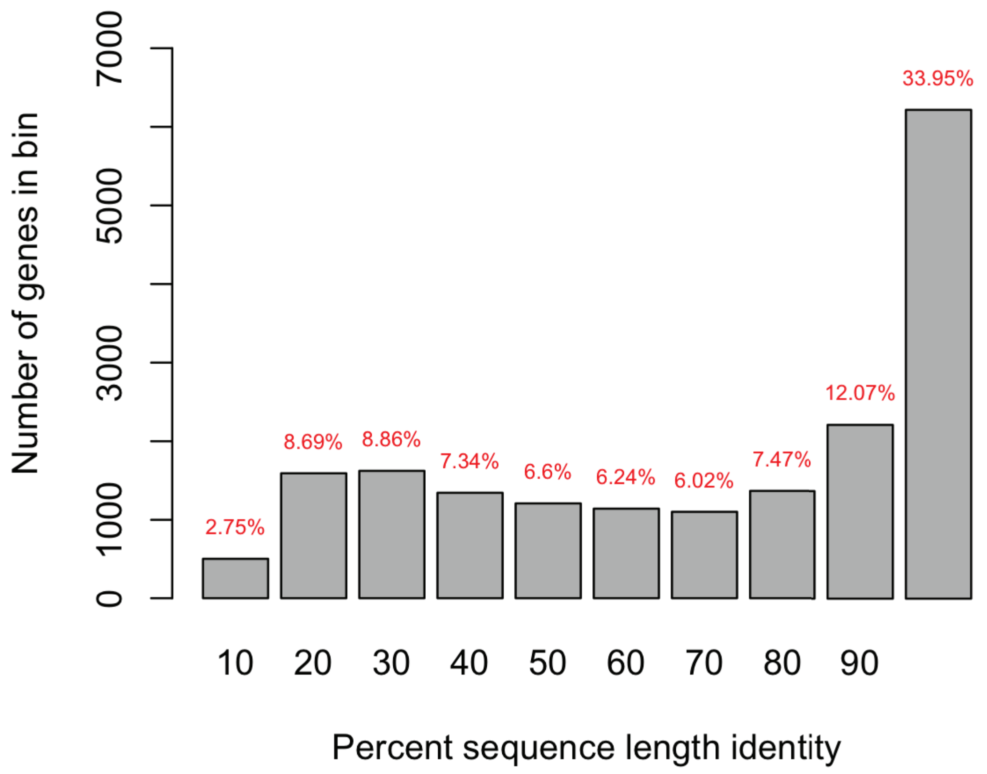


**Figure 1:** Histogram of the number of genes in each percent sequence identity bin. Red text above the bars indicates the percentage of these genes (those with >10% sequence identity) in each bin.

#### 2 Biological replicate validation

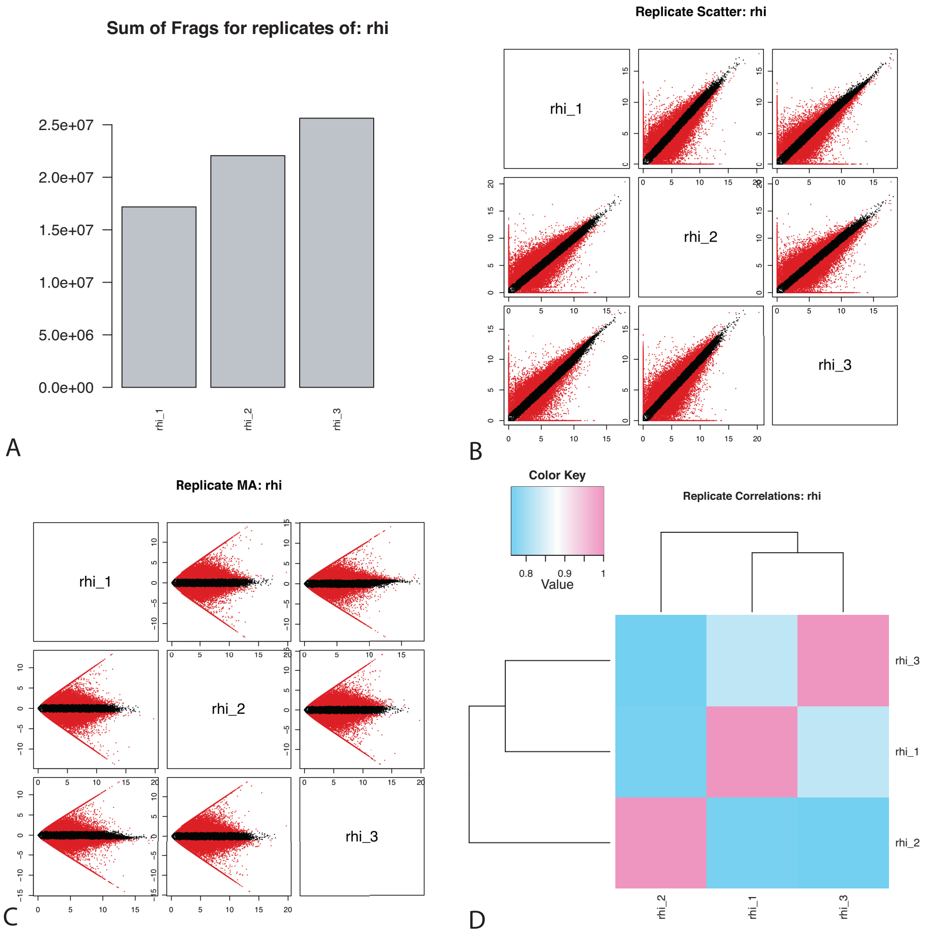


**Figure 2:** Replication information for the three biological replicates of rhizome meristem (RHI) tissue, showing A) the sum of fragments for each replicate, B) 1:1 scatterplots of transcript abundance for all pairwise comparisons of the replicates, C) MA plots (Bland Altman plot) for all pairwise comparisons of the replicates, and D) the correlation coefficients between all replicates.

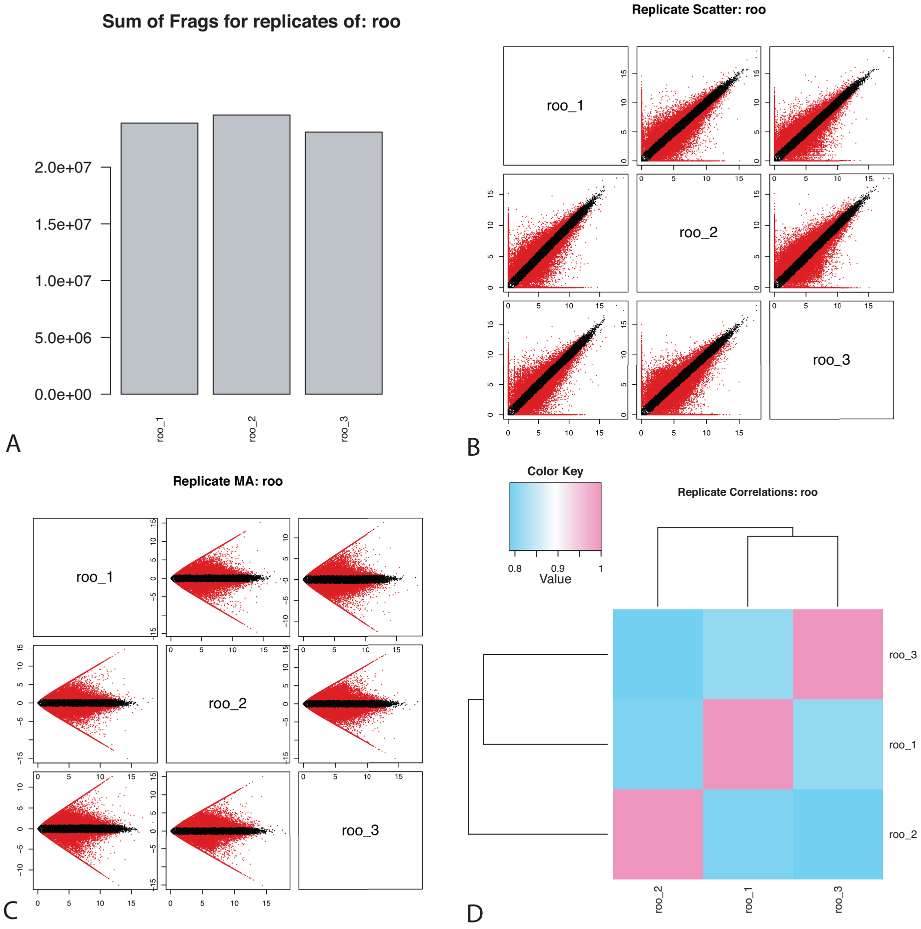


**Figure 3:** Replication information for the three biological replicates of root (ROO) tissue, showing A) the sum of fragments for each replicate, B) 1:1 scatterplots of transcript abundance for all pairwise comparisons of the replicates, C) MA plots (Bland Altman plot) for all pairwise comparisons of the replicates, and D) the correlation coefficients between all replicates.

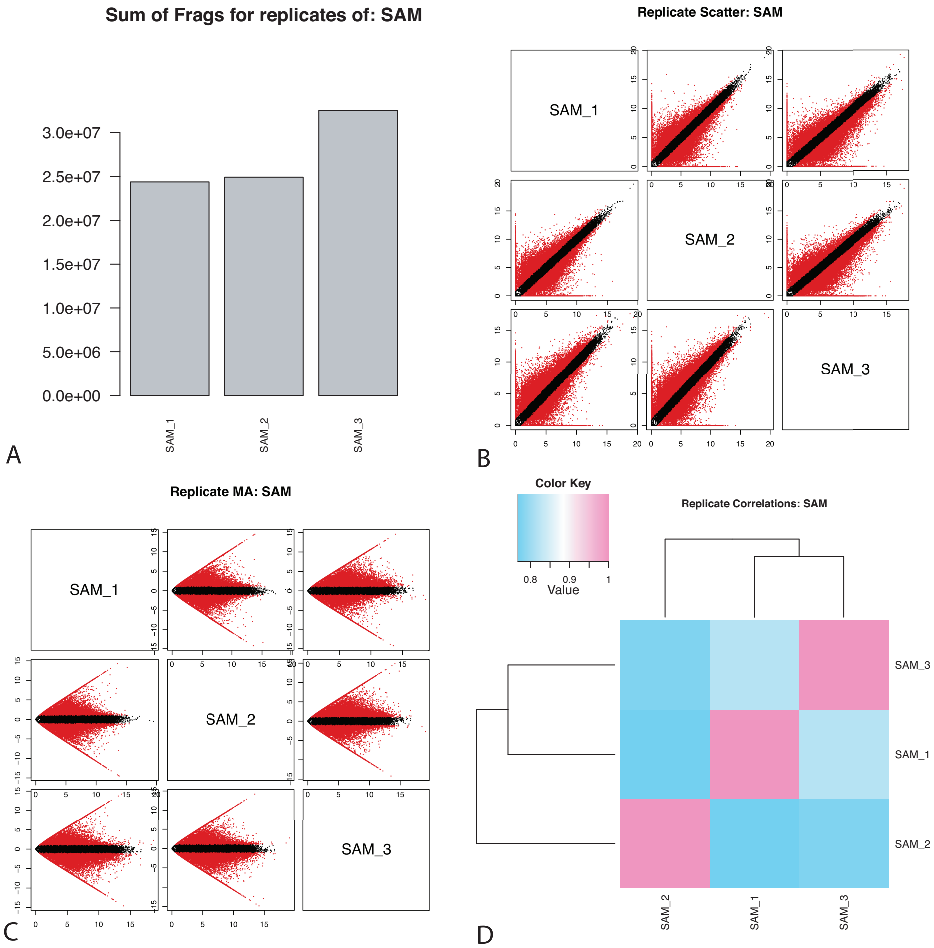


**Figure 4:** Replication information for the three biological replicates of shoot apical meristem (SAM) tissue, showing A) the sum of fragments for each replicate, B) 1:1 scatterplots of transcript abundance for all pairwise comparisons of the replicates, C) MA plots (Bland Altman plot) for all pairwise comparisons of the replicates, and D) the correlation coefficients between all replicates.

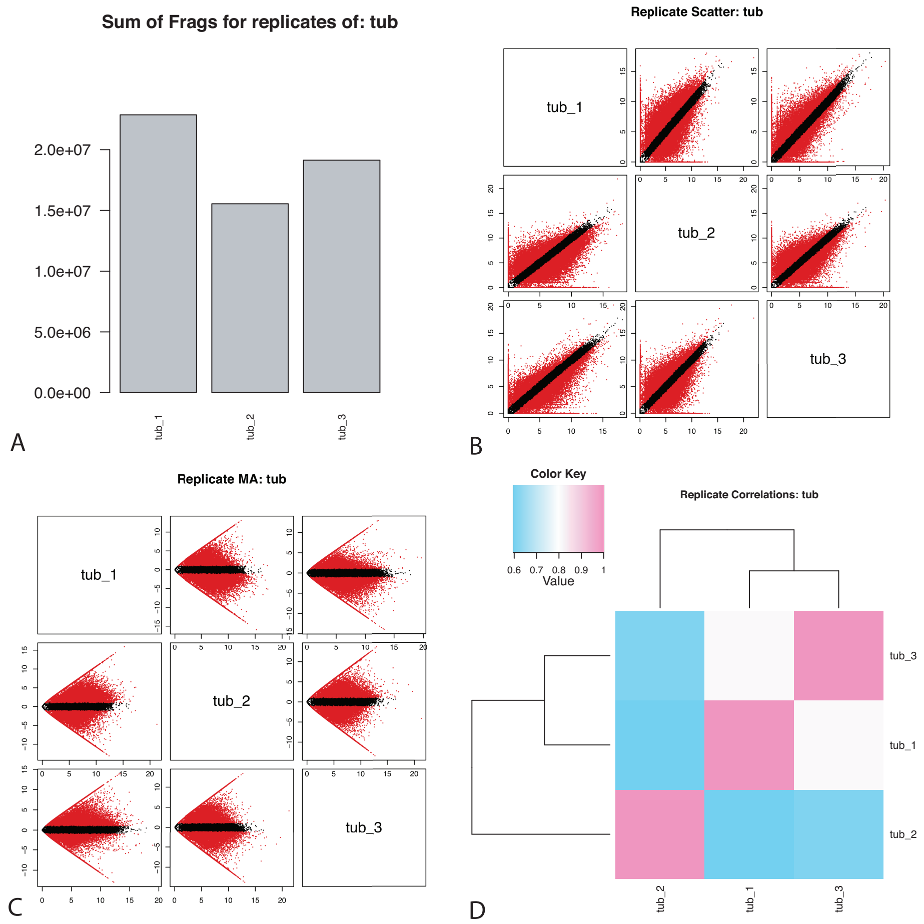


**Figure 5:** Replication information for the three biological replicates of tuberous root (TUB) tissue, showing A) the sum of fragments for each replicate, B) 1:1 scatterplots of transcript abundance for all pairwise comparisons of the replicates, C) MA plots (Bland Altman plot) for all pairwise comparisons of the replicates, and D) the correlation coefficients between all replicates.

### 3 Candidate gene groups

**Table 2:** Search terms used to identify isoforms in candidate gene groups. We queried the annotated transcriptome using R's grep() function to identify the isoforms and then checked their expression levels in the RSEM data using deseq2.

| Gene Group | Search Term | Search Column | Num Isoforms |
|---|---|---|---|
| Starch | GO:0019252 | gene_ontology_blast | 168 |
| Cytokinin | GO:0009735 | gene_ontology_blast | 356 |
| Abscisic Acid | GO:0009737 | gene_ontology_blast | 1357 |
| Auxin | GO:0009733 | gene_ontology_blast | 716 |
| MADS-Box | MADS-box | sprot_Top_BLASTX_hit | 116 |
| KNOX | KNOX | Pfam | 18 |
| Gibberellin | GO:0009739 | gene_ontology_blast | 295 |
| Expansins | Expansin | sprot_Top_BLASTX_hit | 117 |
| Lignin | GO:0009809 | gene_ontology_blast | 325 |
| 14-3-3 genes | 14-3-3-like protein | sprot_Top_BLASTX_hit | 62 |
| CDPK | Calcium-dependent protein kinase | sprot_Top_BLASTX_hit | 180 |
| Lipoxygenase | lipoxygenase | gene_ontology_blast | 54 |

**Table 3:** To identify the expression levels of isoforms corresponding to specific candidate genes, we blasted amino acid sequences of the specific candidate gene to a blast-database of the assembled transcriptome.

| Candidate Gene | Source | Accession Number | Taxon |
|---|---|---|---|
| FT-like | GenBank | AGZ20207.1 | *Allium cepa* |
| FT-like | GenBank | AGZ20210.1 | *Allium cepa* |
| FT-like | SPROT | tr\|M1C558\|M1C558_SOLTU | *Solanum tuberosum* |
| Sulfite reductase | GenBank | AAC24584.1 | *Prunus armeniaca* |
| WOX4 | SPROT | sp\|Q6X7J9\|WOX4_ARATH | *Arabidopsis thaliana* |
| OsbHLH120 | SPROT | tr\|Q67TR8\|Q67TR8_ORYSJ | *Oryza sativa* |
| IDD5 | SPROT | sp\|Q9ZUL3\|IDD5_ARATH | *Arabidopsis thaliana* |

**Table 4:** Gene families, physiological signaling pathways, and biosynthesis pathways identified in the literature as implicated in USO formation or root thickening. Up-regulation corresponds to an increase of the gene in modified organs vs. non-modified tissue or earlier developmental stages. In Group Pattern and DEGs (referring to *Bomarea multiflora*), up vs. down regulation corresponds to expression levels in TR vs. FR, ns corresponds to nonsignificant results, lde corresponds to a general pattern which is less differentially expressed than expected by chance, and none corresponds to no DEGs in the group. In DEGs, the general pattern of identified members of that group

| | | | ***Bomarea multiflora*** | |
|---|---|---|---|---|
| **Gene Group** | **Original Taxa** | **Pattern in literature** | **Group Pattern** | **DEGs** |
| Abscisic Acid | lotus (Yang et al., 2015), sweet potato (Noh et al., 2010; Dong et al., 2019), and potato (Xu et al., 1998) | up, then down | ns | down |
| Auxin | many (Noh et al., 2010; Cheng et al., 2013b; Sojikul et al., 2015; Yang et al., 2015; Xu et al., 2016; Hannapel et al., 2017; Li et al., 2019; Dong et al., 2019; Kolachevskaya et al., 2019) | up | none | none |
| CDPK | cassava (Sojikul et al., 2010), potato (Raíces et al., 2003), and lotus (Cheng et al., 2013b) | up | ns | up |
| Cytokinin | sweet potato (Noh et al., 2010) | up | ns | up and down |
| Expansins | cassava (Sojikul et al., 2015), *Callerya speciosa* (Xu et al., 2016), lotus (Cheng et al., 2013b), Convulvulaceae (Eserman et al., 2018), and potato (Jung et al., 2010) | up | down | none |
| Gibberellin | various roots (Tanimoto, 2012), *Callerya speciosa* (Xu et al., 2016), wild sweet potato (Li et al., 2019), and sweet potato (Dong et al., 2019) | down | none | down |
| Knox | sweet potato (Tanaka et al., 2008) | up | none | none |
| Lignin | cassava (Sojikul et al., 2015), wild sweet potato (Li et al., 2019), and *Callerya speciosa* (Xu et al., 2016) | down | down | down |
| MADS-Box | wild sweet potato (Li et al., 2019), sweet potato (Noh et al., 2010; Dong et al., 2019), lotus (Cheng et al., 2013b), and *Sagittaria trifolia* (Cheng et al., 2013a) | up | up | up |
| Starch | cassava (Sojikul et al., 2010, 2015), lotus (Cheng et al., 2013b; Yang et al., 2015), wild and domestic sweet potato (Eserman et al., 2018; Li et al., 2019; Dong et al., 2019), and potato (Xu et al., 1998) | up | up | up |
| 14-3-3 | *Arabidopsis thaliana* (Van Kleeff et al., 2014; Mayfield et al., 2012; He et al., 2015) | up (in stout roots) | lde | none |
| Lipoxygenases | *Solanum tuberosum* (Kolomiets et al., 2001) *Brassica spp.* (Hearn et al., 2018) | up | lde | none |

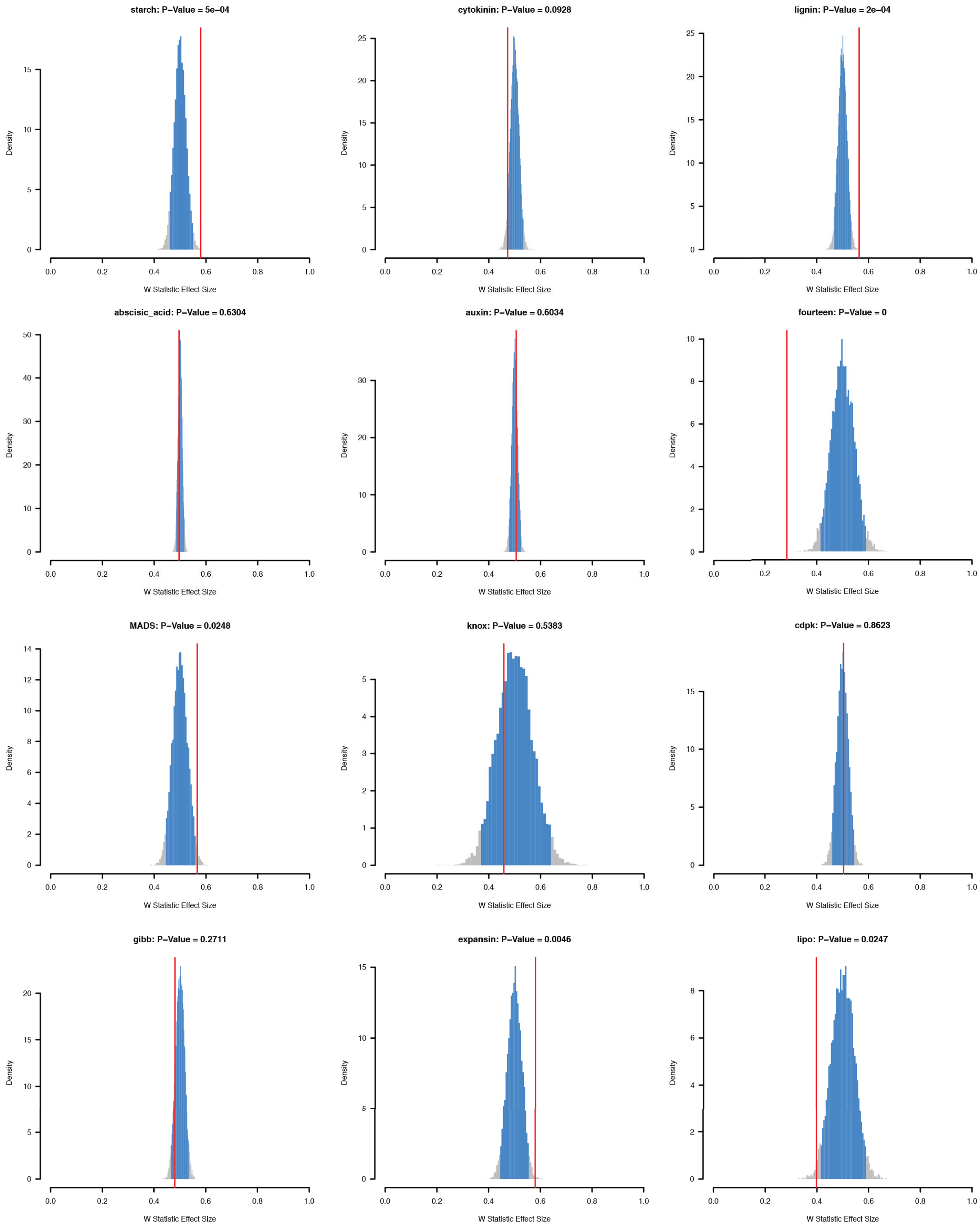


**Figure 6:** For each group of genes, the simulated distribution of effect sizes from a Mann-Whitney U test comparing the absolute values of log2-fold changes of randomly selected groups of n genes to the full pool of genes. Bars in blue represent 95% of the distribution. Grey bars fall outside of the 95% confidence interval. Horizontal red lines indicate the actual effect size observed for the group of genes in question. Red lines that fall to the right of the 95% confidence interval are more differentially expressed than the null expectation, red lines within the interval are non-significant, and red lines to the left of the interval are less differentially expressed than the null expectation.

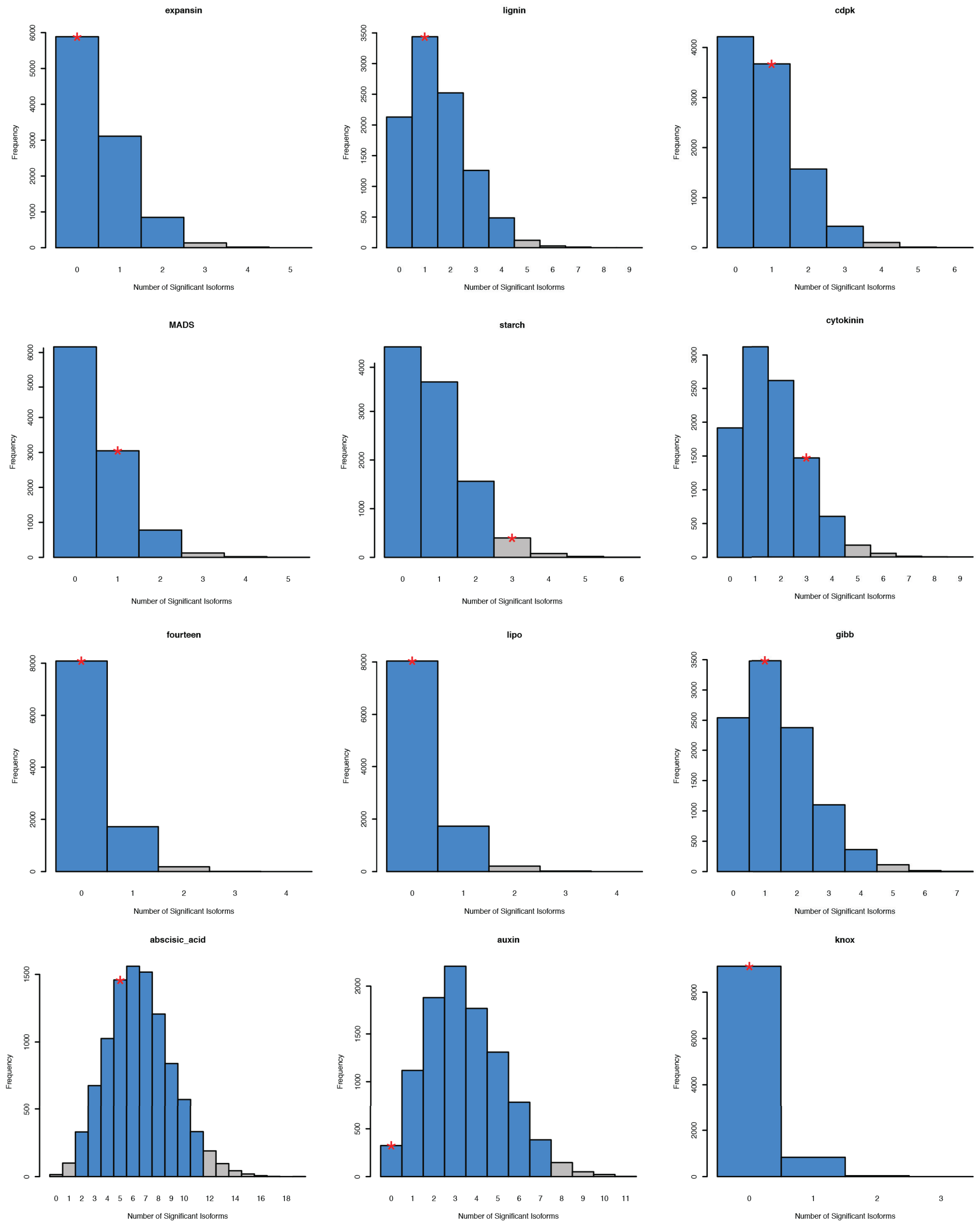


**Figure 7:** For each group of genes, the simulated distribution of expected numbers of DEGs. Bars in blue represent 95% of the distribution. Grey bars fall outside of the 95% credible set. Horizontal red lines indicate the actual number of DEGs observed in the group of genes in question. Red lines that fall to the right of the 95% credible set are more differentially expressed than the null expectation and red lines within the interval are non-significant.

### 4 PEBP Gene Family Evolution

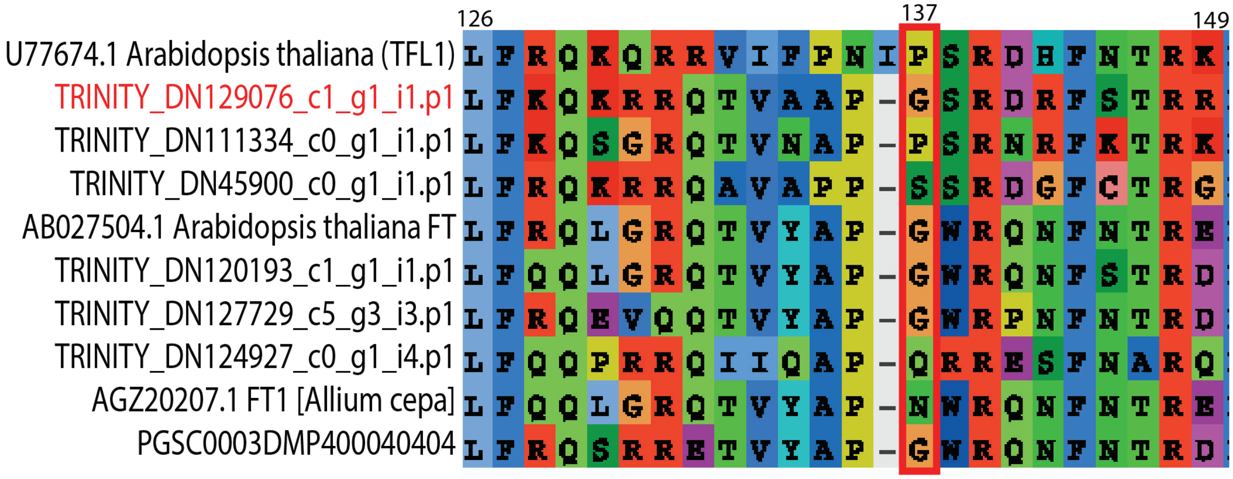


**Figure 8:** Amino acid alignment of *Bomarea multiflora* PEBP orthologs, characterized *Arabidopsis* FT and TFL1 copies, and other PEBP genes implicated in USO development. DEG *Bomarea* copy is labeled in red. AthFT position G137 is outlined in red, showing a glycine mutation in the *Bomarea* TFL DEG.
